# A fast and agnostic method for bacterial genome-wide association studies: bridging the gap between kmers and genetic events

**DOI:** 10.1101/297754

**Authors:** Magali Jaillard, Leandro Lima, Maud Tournoud, Pierre Mahé, Alex van Belkum, Vincent Lacroix, Laurent Jacob

## Abstract

**Motivation:** Genome-wide association study (GWAS) methods applied to bacterial genomes have shown promising results for genetic marker discovery or fine-assessment of marker effect. Recently, alignment-free methods based on kmer composition have proven their ability to explore the accessory genome. However, they lead to redundant descriptions and results which are hard to interpret.

**Methods:** Here, we introduce DBGWAS, an extended kmer-based GWAS method producing interpretable genetic variants associated with phenotypes. Relying on compacted De Bruijn graphs (cDBG), our method gathers cDBG nodes identified by the association model into subgraphs defined from their neighbourhood in the initial cDBG. DBGWAS is fast, alignment-free and only requires a set of contigs and phenotypes. It produces annotated subgraphs representing local polymorphisms as well as mobile genetic elements (MGE) and offers a graphical framework to interpret GWAS results.

**Results:** We validated our method using antibiotic resistance phenotypes for three bacterial species. DBGWAS recovered known resistance determinants such as mutations in core genes in *Mycobacterium tuberculosis* and genes acquired by horizontal transfer in *Staphylococcus aureus* and *Pseudomonas aeruginosa* – along with their MGE context. It also enabled us to formulate new hypotheses involving genetic variants not yet described in the antibiotic resistance literature.

**Conclusion:** Our novel method proved its efficiency to retrieve any type of phenotype-associated genetic variant without prior knowledge. All experiments were computed in less than two hours and produced a compact set of meaningful subgraphs, thereby outperforming other GWAS approaches and facilitating the interpretation of the results.

**Availability:** Open-source tool available at https://gitlab.com/leoisl/dbgwas

## Introduction

The aim of Genome-Wide Association Studies (GWAS) is to identify associations between genetic variants and a phenotype observed in a population. They have recently emerged as an important tool in the study of bacteria, given the availability of large panels of bacterial genomes combined with phenotypic data (Farhat et al., 2013; Sheppard et al., 2013; Alam et al., 2014; Chewapreecha et al., 2014; Earle et al., 2016; Lees et al., 2016; Jaillard et al., 2017b).

GWAS require encoding the genomic variation as numerical factors. The most common approaches rely on single nucleotide polymorphisms (SNPs), defined by aligning all genomes in the panel against a reference genome (Farhat et al., 2013; Alam et al., 2014; Chewapreecha et al., 2014) and on gene presence/absence, using a pre-defined collection of genes (Earle et al., 2016; Jaillard et al., 2017b). Relying on SNPs or gene presence/absence is reasonable when studying species whose genomic variations can be summarised by a list of pre-defined biological entities. However, a suitable reference is not always available for bacteria, particularly for species with a large accessory genome – the part of the genome which is not present in all strains. Moreover, when focusing on the variation in gene content, one would be unable to cover variants in noncoding regions, including those related to transcriptional and translational regulation (Zhang et al., 2013; Blair et al., 2015).

To circumvent these issues and make bacterial genomes amenable to GWAS, recent studies have relied on kmers: all nucleotide substrings of length *k* found in the genomes (Sheppard et al., 2013; Earle et al., 2016; Lees et al., 2016). Kmers enable to account for diverse genetic events such as the acquisition of SNPs, (long) insertions/deletions and recombinations. Unlike SNP- or gene-based approaches, kmer-based approaches do not require a reference genome or any assumption on the nature of the causal variants and can even be performed without having to assemble the genome sequences (Le Bras et al., 2016).

While kmers can reflect any genomic variation in a panel, they do not themselves represent biological entities. Translating the result of a kmer-based GWAS into meaningful genetic variants typically requires mapping a large and redundant set of short sequences (Sheppard et al., 2013; Earle et al., 2016; Lees et al., 2016; Rahman et al., 2017). Recent studies have suggested reassembling the significantly associated kmers to reduce redundancy and retrieve longer sequences (Lees et al., 2016; Rahman et al., 2017). Nonetheless, kmer representation often loses in interpretability what it gains in flexibility, and the best way to encode the genomic variation in bacterial GWAS is not yet clearly defined (Read and Massey, 2014; Power et al., 2017).

Our approach, coined DBGWAS, for *De Bruijn Graph GWAS*, bridges the gap between, on the one hand, SNP- and gene-based representations lacking the right level of flexibility to cover complete genomic variation, and, on the other hand, kmer-based representations which are flexible but not readily interpretable. We use De Bruijn graphs (de Bruijn, 1946) (DBGs), which are widely used for *de novo* genome assembly (Pevzner et al., 2001; Zhang et al., 2011) and variant calling (Iqbal et al., 2012; Le Bras et al., 2016). These graphs connect overlapping kmers (here DNA fragments), yielding a compact summary of all variations across a set of genomes. Figure 1 illustrates the construction of such a graph for a simple example, where the only variation among the aligned genomes is a point mutation. DBGs also accommodate more complex disparities including rearrangements and insertions/deletions (Supplementary Figure S1).

**Figure 1:**
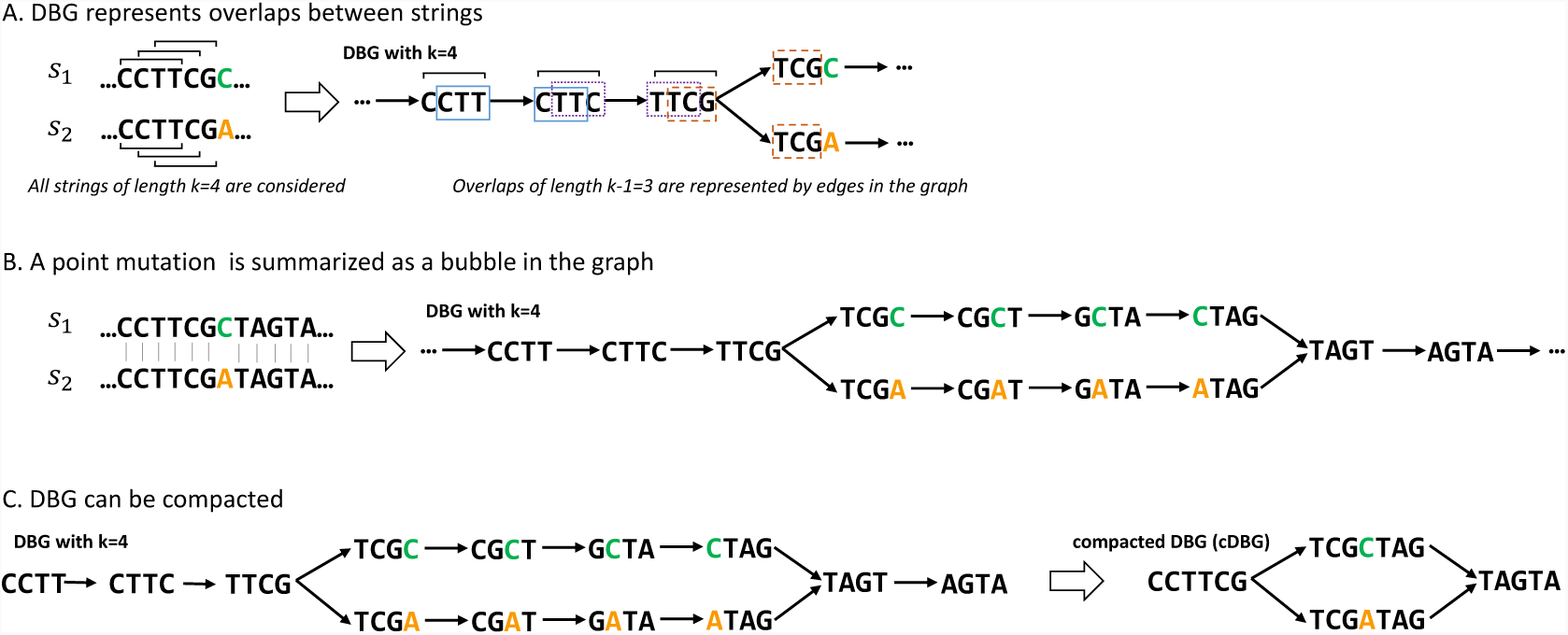
Compacted DBG construction over a set of sequences differing by a single point mutation. In this example two sequences *s*1 and *s*2 of length 12 differ by a single letter. All kmers (*k* = 4) present in these sequences are listed. A) A link is drawn between two kmers when the *k* − 1 = 3 last nucleotides of the first kmer equal the 3 first nucleotides of the second kmer. B) The bubble pattern represents the SNP C to A; each branch of the bubble represents an allele. C) linear paths of the graph are compacted; the compacted DBG of the example only contains four nodes (unitigs) and represents the same variation as the original DBG, which contained 13 nodes (kmers).

DBGWAS relies on the ability of compacted DBGs (cDBGs) to eliminate local redundancy, reflect genome variations, and characterise the genomic environment of a kmer at the population level. More precisely, we build a single cDBG from all the genomes included in the association study (in practice, up to thousands). The graph nodes – called unitigs – represent, by construction, sequences of variable length and are at the right level of resolution for the set of genomes considered, taking into account adaptively the genomic variation. The unitigs are individually tested for association with the phenotype, while controlling for population structure. The unitigs found to be phenotype-associated are then localised in the cDBG. Subgraphs induced by their genomic environment are extracted. They often provide a direct interpretation in terms of genetic events which results from the integration of three types of information: 1) the *topology* of the subgraph, reflecting the nature of the genetic variant, 2) the *metadata* represented by node size and colour, allowing us to identify which unitigs in the subgraph are associated to a particular phenotype status, and 3) an optional sequence *annotation* helping to detect unitig mapping to – or near – a known gene.

We benchmarked our novel method using several antibiotic resistance phenotypes within three bacterial species of various degrees of genome plasticity: *Mycobacterium tuberculosis*, *Staphylococcus aureus* and *Pseudomonas aeruginosa*. The subgraphs built from significant unitigs described SNPs or insertions/deletions in both core and accessory regions and were consistent with results obtained with a targeted resistome-based GWAS approach. However, novel genotype-to-phenotype associations were also suggested.

## Results

DBGWAS generated a set of ordered subgraphs for every panel of microbial strains and tested antibiotics. It computed the q-values for all the unitigs and ordered the subgraphs according to the smallest of their unitig q-value, denoted as min_*q*_. The top subgraphs therefore represented the genomic environment of the unitigs most significantly associated with the tested phenotype, as discussed in Section *step 3* of the Methods section.

The subgraphs we describe below were obtained with DBGWAS using default parameters plus the annotation option. DBGWAS was only provided with contigs and their related phenotypes and did not use any prior information as to the nature or location of potential causal variants. Each run on the three tested species only took between 16 min and 90 min on a single core and required less than 12Gb of memory (Supplementary Table S1).

A synthetic description of the subgraphs discussed in the results is provided in Table 1, while a description of the top subgraphs obtained for all tested antibiotics, is provided in Supplementary Tables S3 to S5. The subgraphs themselves are available at http://leoisl.gitlab.io/DBGWAS_support/experiments.

**Table 1:**
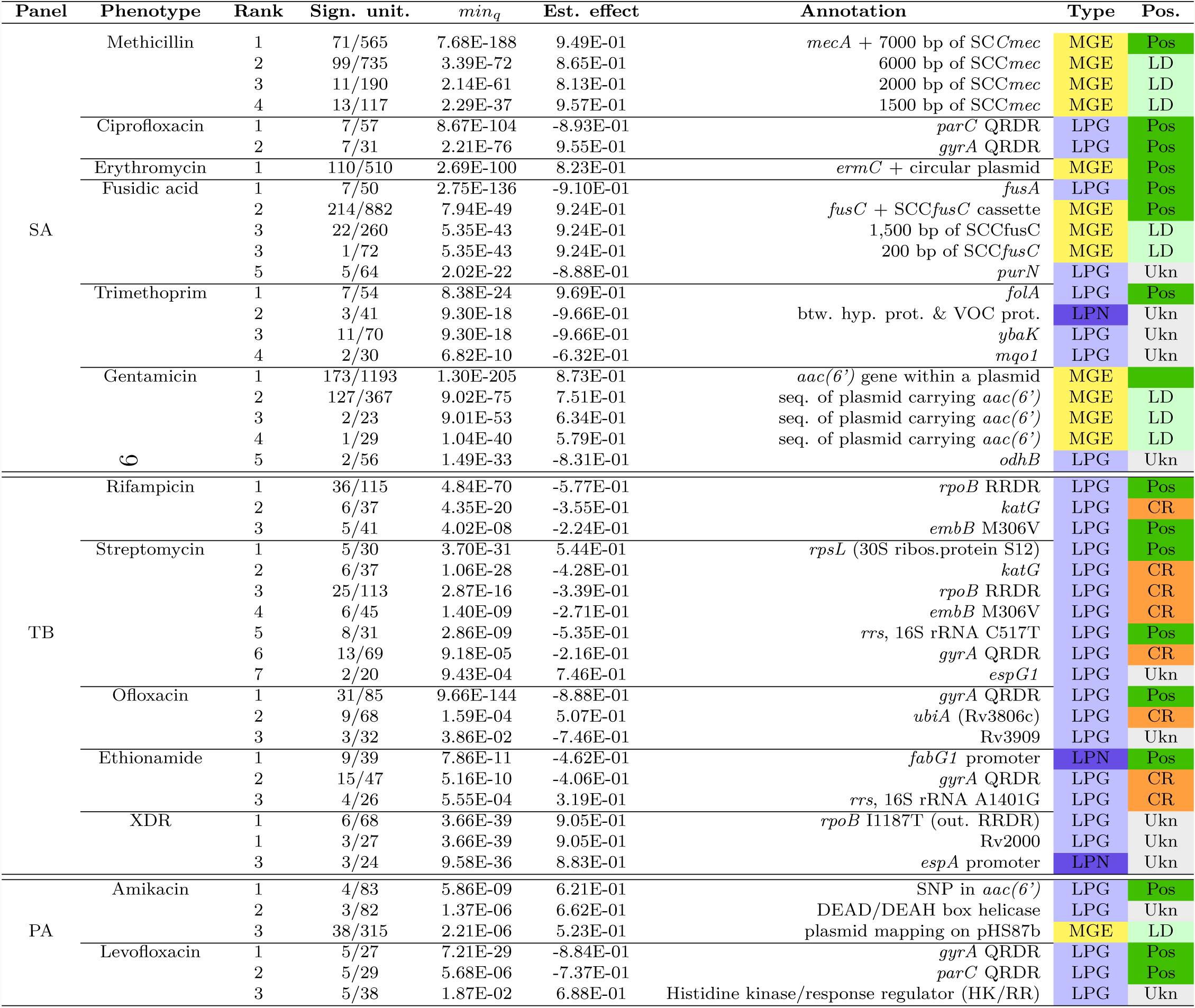
**Resistance determinants identified by DBGWAS for *S. aureus* (SA), *M. tuberculosis* (TB) and *P. aeruginosa* (PA) panels.** For each antibiotic, subgraphs were reported with their rank, number of significant unitigs over all unitigs in the subgraph (Sign. unit.), q-value of the unitig with the lowest q-value (min_*q*_), the corresponding estimated effect (*β* coefficient of the linear mixed model) and annotation of the subgraph. The type of event represented by the subgraph was colour-coded as: yellow for MGE, light blue for local polymorphism in gene (LPG), and dark blue for local polymorphism in noncoding region (LPN). Known positives were indicated in dark green (Pos), regions in LD with a positive in light green (LD), determinants caused by cross-resistance in orange (CR) and unknown determinants in grey (Ukn).

### Coloured bubbles highlight local polymorphism in core genes, accessory genes and noncoding regions

For *P. aeruginosa* levofloxacin resistance, the subgraph obtained with the lowest min_*q*_ highlighted a polymorphic region in a core gene (Figure 2A). Indeed, it showed a linear structure containing a complex bubble, with a fork separating susceptible (blue) and resistant (red) strains. The annotation revealed that all unitigs in this subgraph mapped to the quinolone resistance-determining region (QRDR) of the *gyrA* gene. *gyrA* codes for a subunit of the DNA gyrase targeted by quinolone antibiotics such as levofloxacin and its alteration is therefore a prevalent and efficient mechanism of resistance (Hooper and Jacoby, 2015; Lowy, 2003). In all our experiments related to quinolone resistance, DBGWAS identified QRDR mutations in either *gyrA* or *parC*, which codes for another well-known quinolone target: *P. aeruginosa* levofloxacin (first subgraph, *gyrA*: min_*q*_ = 7.21 × 10^−29^ and second, *parC*: 5.68 × 10^−06^), *S. aureus* ciprofloxacin (first, *parC*: min_*q*_ = 8.67 × 10^−104^ and second, *gyrA*: 2.21 × 10^−76^), and ofloxacin resistance in *M. tuberculosis*, whose genome does not contain the *parC* gene (Piton et al., 2010) (first, *gyrA*: min_*q*_ = 9.66 × 10^−144^).

**Figure 2:**
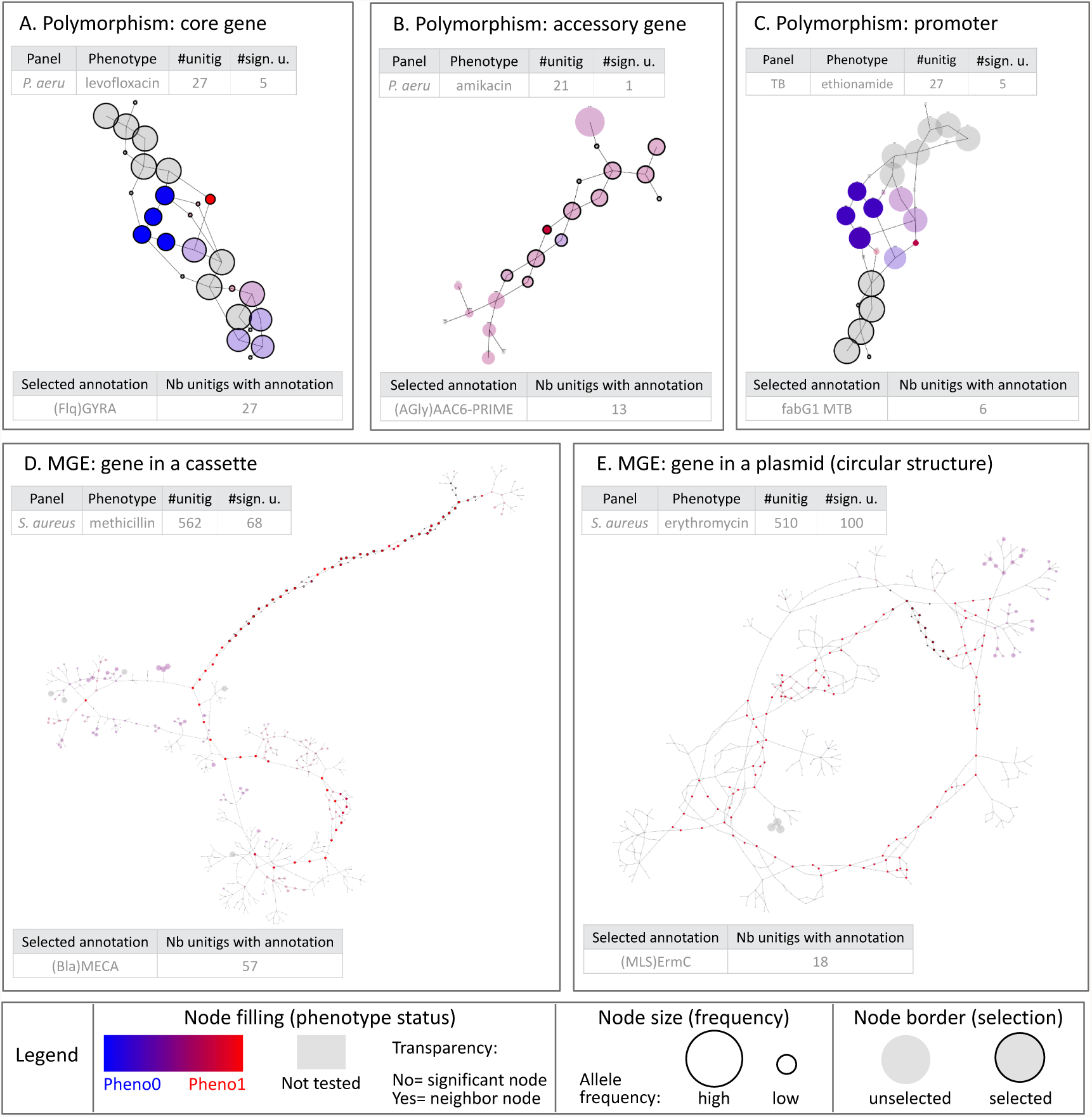
Different types of genetic events identified by DBGWAS. Each subgraph represents a distinct genetic event. Panel A shows the subgraph with lowest min_*q*_ extracted for *P. aeruginosa* levofloxacin resistance. It was composed of 27 unitigs, 5 of which were significantly associated with resistance. Susceptible unitigs are shown in blue, while resistant unitigs in red. All unitigs of this subgraph mapped to the *gyrA* gene. Panels B, C, D, E correspond to the top subgraphs obtained for other panels/phenotypes. The larger the node, the higher the allele frequency. Grey nodes were present in *>* 99% or *<* 1% of the strains and were not tested. Bright blue (resp. bright red) nodes were present almost exclusively in susceptible (resp. resistant) strains. Pale blue (resp. pale red) nodes were present with a larger frequency in susceptible (resp. resistant) strains. Circled black nodes mapped to annotated genes.

For *P. aeruginosa* amikacin resistance, the top subgraph (min_*q*_ = 5.86 × 10^−9^) highlighted a SNP in an accessory gene (Figure 2B). As in Figure 2A, it contained a fork separating a blue and a red node. However, other remaining nodes were not grey: they represented an accessory sequence because they were not present in all the strains. Most of these nodes were pale-red, showing that the accessory sequence was more frequent in resistant samples. The annotation revealed that this subgraph corresponded to *aac(6’)*, a gene coding for an aminoglycoside 6-acetyltransferase, an enzyme capable of inactivating aminoglycosides, such as amikacin, by acetylation (Lambert, 2002). Most unitigs in this gene had a low association with resistance, except for the ones describing this particular SNP. This mutation, L83S, lying in the enzyme binding site, was previously shown to be responsible for substrate specificity alteration in a strain of *Pseudomonas fluorescens* (Lambert et al., 1994). It appeared thus to increase the amikacin acetylation ability of *aac(6’)*, making its association to amikacin resistance more significant than the gene presence itself.

Finally, for *M. tuberculosis* ethionamide resistance, the top subgraph (min_*q*_ = 7.86 × 10^−11^, Figure 2C) represented a polymorphic region in a core gene promoter. The subgraph was mostly grey and linear with a localised blue and red fork. The most reliable annotation for this subgraph was *fabG1* (also known as *mabA*), a core gene previously shown to be involved in ethionamide and isoniazid resistance (Lee et al., 2000; Farhat et al., 2016). None of the significantly associated unitigs mapped to the *fabG1* gene, but their close neighbours did (highlighted in Figure 2C by black circles), suggesting that the detected variant was located in the promoter region of the gene. This was confirmed by mapping the significant unitig sequences using the Tuberculosis Mutation database of the *mubii* resource (Flandrois et al., 2014).

### Long single-coloured paths denote mobile genetic element insertions

For *S. aureus* resistance to methicillin, the top subgraph (min_*q*_ = 7.68 × 10^−188^), shown in Figure 2D, revealed a gene cassette insertion. It contained a long path of red nodes, and a branching region including another red node path. The first path mapped to the *mecA* gene, extensively described in this context and known to be carried by the Staphylococcal Cassette Chromosome *mec* (SCC*mec*) (Lowy, 2003; IWG-SCC consortium, 2009; Gordon et al., 2014). The other part of the subgraph represented a *>*5,000 bp fragment of the cassette. It was less linear because it summarised several types of the cassette differing by their structure and gene content (IWG-SCC consortium, 2009). The next subgraphs represented other regions of the same cassette. Interestingly, considering a greater number of unitigs to build the subgraphs would lead to merging these individual subgraphs, representing related genomic regions, into a single subgraph. This can be done by increasing the Significant Features Filter (*SFF*) parameter value which defines the unitigs used to build the subgraphs. By default, the unitigs corresponding to the 100 lowest q-values are retained (*SFF* = 100). Increasing the *SFF* value to 150 (150th q-value = 1.60 × 10^−27^) allowed us to reconstruct the entire SCC*mec* cassette, as shown in Supplementary Figure S3.

For *S. aureus* erythromycin resistance, a unique subgraph was generated (min_*q*_ = 2.69 × 10^−100^). As shown in Figure 2E, the subgraph described the circular structure of a 2,500 bp-long plasmid known to carry the causal *ermC* gene (Westh et al., 1995; Gordon et al., 2014) together with a replication and maintenance protein in strong linkage disequilibrium with *ermC*.

For *P. aeruginosa* amikacin resistance, the third subgraph (min_*q*_ = 2.21 × 10^−6^) represented a 10,000 bp plasmid acquisition. Using the NCBI nucleotide database (Benson et al., 2012), most of the unitigs in this subgraph mapped to the predicted prophage regions of an integrative and conjugative plasmid, whose structure was recently described as the pHS87b plasmid in the amikacin resistant *P. aeruginosa* HS87 strain (Bi et al., 2016). Supplementary Figures S4 and S5 provide more examples of MGEs recovered by DBGWAS, and Section *step 3* of the Methods discusses *SFF* default value and tuning.

### Comparison of DBGWAS to reference- and kmer-based methods: DBGWAS reports expected variants without prior knowledge, with the highest computational efficiency

DBGWAS relies on bugwas (Earle et al., 2016) – a state-of-the-art association model for bacterial GWAS – to test for significant associations between unitigs and phenotypes. The performance of detecting true associations using unitigs was previously assessed using simulated data (Jaillard et al., 2017a). In this preliminary study, we showed that the linear mixed model implemented by bugwas presented the best power to detect genuine associations under different population structure hypotheses, among several association models.

Here, we evaluated DBGWAS using real data. Although resistance determinants are not perfectly and exhaustively known in any species, some resistance mechanisms are well described enough to allow evaluation on real data. This is the case of target alteration in fluoroquinolone resistance or, in *M. tuberculosis* resistance, to antibiotics of the aminoglycoside family. We thus compared resistance determinants obtained by DBGWAS for *M. tuberculosis* (aminoglycoside) streptomycin resistance and *P. aeruginosa* (fluoroquinolone) levofloxacin resistance, to determinants obtained by a resistome-based GWAS (RWAS) strategy (Davis et al., 2016; Jaillard et al., 2017b), as described in the Methods section, and by two other recent kmer-based methods: SEER (Lees et al., 2016) and HAWK (Rahman et al., 2017). For *P. aeruginosa* levofloxacin resistance (Figure 3A), DBGWAS and SEER found both known causal determinants reported by the RWAS strategy, *gyrA* and *parC*, while HAWK only reported *gyrA*. SEER reported 403 kmers, all linked to *gyrA* and *parC* contrary to others methods that all reported less than 10 features, among which new hypotheses. For *M. tuberculosis* streptomycin resistance (Figure 3B), the four methods reported both known causal determinants *rpsL* and *rrs*, however not always in the same order. Indeed, while the RWAS and DBGWAS methods found the causal *rpsL* determinant as the first position, SEER and HAWK reported first the *katG* determinant. All the methods identified several markers described for other antibiotics. This observed cross-resistance to antibiotics is a well described phenomenon in *M. tuberculosis* species (Traore et al., 2000; Palomino and Martin, 2014). Compared to SEER and HAWK, DBGWAS produced a smaller number of features (24 *versus* several thousands), in a shorter time (1h 18m *versus >*9h), without loss of sensitivity regarding the detection of resistance markers. Additional results for all the antibiotics can be found in Supplementary Tables S6 and S7 for RWAS, and in Supplementary Tables S3 and S5 for DBGWAS.

**Figure 3:**
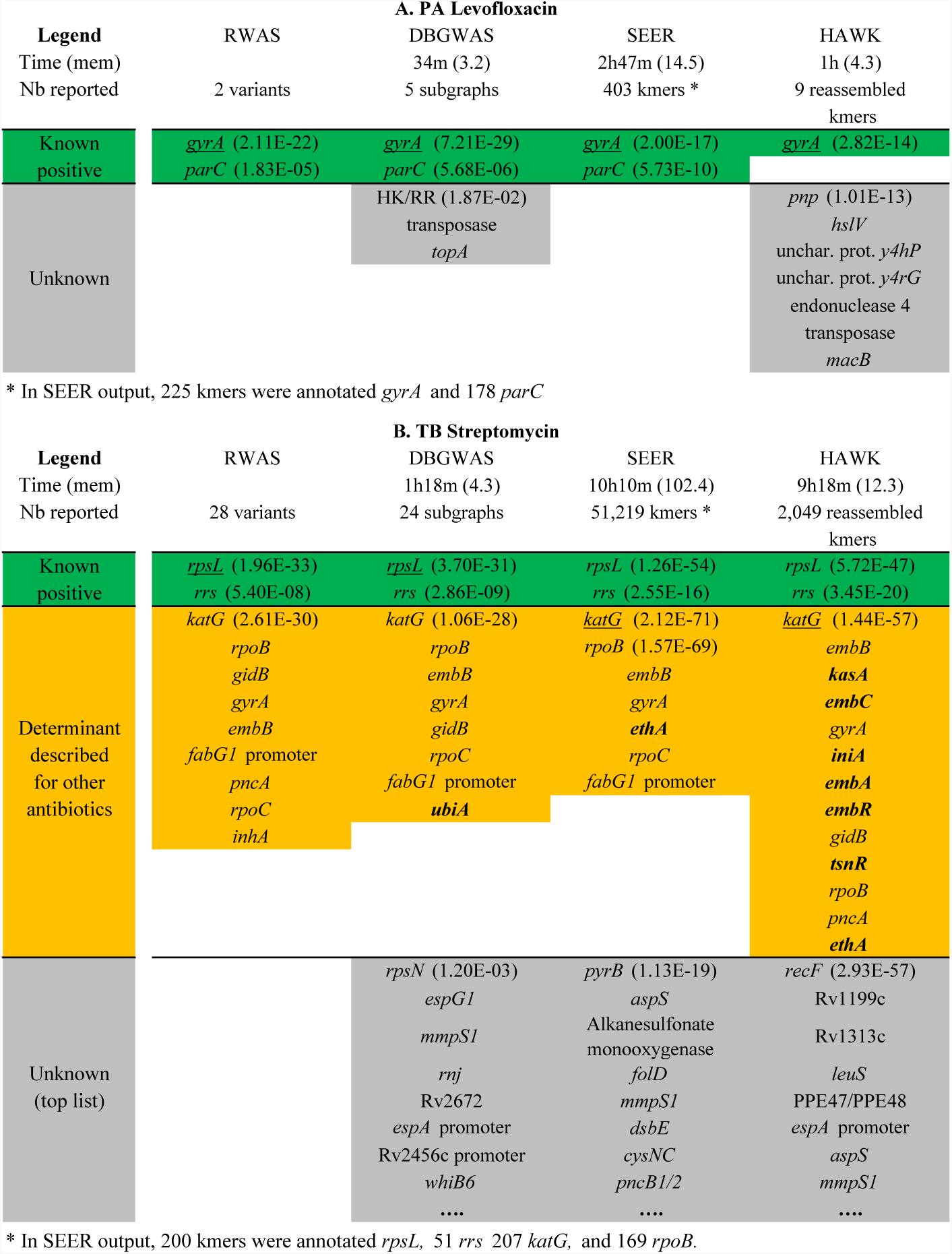
Resistance determinants found by the 4 methods, for *M. tuberculosis* streptomycin and *P. aeruginosa* levofloxacin resistances. In this figure, we report deduplicated annotations of features identified as significant with the default parameters (p-value for SEER and HAWK or q-value for RWAS and DBGWAS). The total number of reported features is given in the header. For kmer-based methods, annotations were retrieved by mapping unitig/kmer sequences on the resistance and Uniprot databases. Green cells correspond to resistance determinants already described in the literature, orange cells to resistance determinants described for association with other antibiotics (annotations not found by RWAS are written in bold), and grey cells to unknown determinants. Within each category, annotations are ordered by increasing minimum p/q-values, corresponding to the lowest p/q-value found for each annotation before deduplication (p/q-values are reported only for the most significant annotations). For each method, the annotation with the lowest p/q-values is underlined. The execution time and memory load (in Gigabytes) are shown in the header (see also Supplementary Table S2).

In addition to resistance markers, the three kmer-based approaches reported several unknown determinants, not described in the context of resistance. Within them, in the context of streptomycin resistance, the *mmpS1* annotation was reported by the three methods, but not by the RWAS approach, as this gene was not included in the targeted approach prior. More generally, any reference-based approaches such as SNP- or gene-based GWAS or RWAS are limited in the context of new marker discovery, especially for species with a large accessory genome, since any causal variant absent from the chosen reference would remain non-tested. Besides being time-consuming, preparing such a list of genetic variants can even be problematic for bacterial species without extensive annotation nor reference availability.

Agnostic approaches avoid the difficulty of designing an exhaustive variant database for the GWAS. However, HAWK and SEER reported several thousands kmers for *M. tuberculosis* streptomycin resistance, while DBGWAS reported only 24 annotated subgraphs without missing expected determinants (Figure 3A). Indeed, when several phenotype-associated unitigs were found within a particular region of the genome, DBGWAS gathered them into a single subgraph enriched with metadata and annotation (Supplementary Section 6), providing a valuable interpretation framework. As an example, the top subgraph for rifampicin resistance (min_*q*_ = 4.84×10^−70^) contained 36 significant unitigs, either blue or red. Instead of a single point mutation, this subgraph represented a polymorphic region known as the rifampicin resistance-determining region (RRDR) of the *rpoB* gene. The unitig with the lowest q-value covered several mutant positions, defining a haplotype strongly associated with rifampicin resistance. Where DBGWAS reported in this case only one subgraph, SEER, for instance, reported 470 kmers with the *rpoB* annotation.

Finally, DBGWAS took less than 2 hours in all our experiments, while SEER took more than one week in some experiments, and HAWK usually ran in less than one day but failed on the most complex dataset composed of genomes of different species. Moreover, SEER required much more memory (up to 100Gb) than DBGWAS and HAWK (Supplementary Table S2).

### DBGWAS suggests novel hypotheses

As DBGWAS screens the genomic variations without prior knowledge, it documented associations never previously described in resistance literature.

In our *P. aeruginosa* panel, the second subgraph obtained for amikacin resistance (min_*q*_ = 1.37 × 10^−6^) gathered unitigs mapping to the 3’ region of a DEAD/DEAH box helicase known to be involved in stress tolerance in *P. aeruginosa* (Illakkiam et al., 2014). The unitig with the lowest q-value was present in 13 of 47 resistant strains and in only 1 of 233 susceptible strains and represented a C-C haplotype summarising two mutated positions: 2097 and 2103. In *P. aeruginosa* levofloxacin resistance, the third subgraph (min_*q*_ = 1.87 × 10^−2^) represented a L650M amino-acid change in a hybrid sensor histidine kinase/response regulator. Such two-components regulatory systems play important roles in the adaptation of organisms to their environment, for instance in the regulation of biofilm formation in *P. aeruginosa* (Ali-Ahmad et al., 2017), and as such may play a role in antibiotic resistance.

In *S. aureus*, polymorphisms within genes not known to be related to resistance were identified for several antibiotics: *purN* (min_*q*_ = 2.02×10^−22^) for fusidic acid, *odhB* (min_*q*_ = 1.49 × 10^−33^) for gentamicin, *ybaK* and *mqo1* (min_*q*_ = 9.30 × 10^−18^, resp. 6.82 × 10^−10^) for trimethoprim. None of these genes have been associated with antibiotic resistance before, to the best of our knowledge.

In *M. tuberculosis*, polymorphisms in two genes encoding proteins involved in *cell wall and cell processes*, *espG1* and *espA*, were found associated with streptomycin (seventh subgraph, min_*q*_ = 9.43×10^−4^) and XDR phenotype (third subgraph, min_*q*_ = 9.58 × 10^−36^) respectively. Again, these genes have never been reported in association with antibiotic resistance before.

Although experimental validation would be required to tell whether these hypotheses are false positive (e.g., in linkage with causal variants) or actual resistance mechanisms not yet documented, DBGWAS is a valuable tool for novel candidate screening. Moreover it provides a first level of variant description (SNPs in gene or promoter, MGE, etc.) which can directly drive the biological validation.

## Discussion

In this article we introduce an efficient method for bacterial GWAS. Our method is agnostic: it screens all genomic variations and is able to identify potential new causal variants as different as SNPs or (MGE) insertions/deletions. It performs as well as the current SNP- and gene-based gold standard approaches for retrieving known determinants, while these standard approaches require strong prior assumptions often limiting the variant search space and requiring fastidious preprocessing.

Our original method, exploiting the genetic environment of the significant kmers, through their neighbourhood in the cDBG, provides a valuable interpretation framework. Because it uses only contig sequences as input, it allows GWAS on bacterial species for which the genomes are still poorly annotated or lack a suitable reference genome. Our method, DBGWAS, makes bacterial GWAS possible in less than two hours using a desktop computer, outperforming state-of-the-art kmer-based approaches.

Underlying our method, graph-based genome sequence representations such as DBGs, extend the notion of the reference genome to cases where a single sequence stops being an appropriate approximation (Marschall et al., 2016; Paten et al., 2017). As demonstrated in this paper, they pave the way to GWAS on highly plastic bacterial genomes and would also be useful for microbiomes (Baaijens et al., 2017) or human tumours (Rahman et al., 2017).

DBGWAS could be extended to different statistical tasks by adapting its underlying association model, to allow for continuous phenotypes or identifying epistatic effects, for instance. The interpretability of the extracted subgraphs could also be improved by training a machine learning model to predict which types of event they represent. This automated labelling could guide users in their interpretation and allow them to search for specific events, such as SNPs in core genes or rearrangements. Knowing the type of event that a subgraph represents could also be of use for constructing a method controlling false discovery rate at the genetic event level (SNPs, MGE insertion) instead of at the unitig level.

A variety of current studies describes computerised models for defining a genomic antibiogram and hopes are high that such technologies will replace the classical methods. Extensive studies have been performed for a multitude of organisms and the more clonal the bacterial species, the more direct homology searches for resistance genes become reliable (Dunne Jr et al., 2017). Several studies have already demonstrated that genomic antibiograms are at least as good as classic phenotypic ones (Gordon et al., 2014). Contrary to our approach, these studies require extensive resistance marker databases. DBGWAS will surely contribute to the extension of such databases or to the development of agnostic genomic antibiograms.

In conclusion, we demonstrate for three medically important bacterial species that resistance markers can be detected rapidly with relative ease, using simple computer equipment. New links between genomic variations and phenotypes can be inferred, providing our method with a clear advantage in comparison to existing procedures. Using our graphical interface will provide future users in all domains of microbiology with an enhanced insight into genotype to phenotype correlation, also beyond antibiotic resistance. This will include complex traits such as biofilm formation, epidemicity and virulence.

## Methods

### Encoding genomic variation with compacted DBGs

DBGs are directed graphs that efficiently represent all the information contained in a set of sequences. Nodes represent all the unique kmers (genome sequence substrings of length *k*) extracted from the input sequences. Edges represent (*k −* 1)-exact-overlaps between kmers: an edge connects a node *n*_1_ to a node *n*_2_ if and only if the (*k −* 1)-length-suffix of *n*_1_ equals the (*k* − 1)-length-prefix of *n*_2_ (Figure 1A).

These graphs can be compacted into cDBGs by merging linear paths (sequences of nodes not linked to more than two other nodes) into a single node referred to as a *unitig* (Butler et al., 2008; Zerbino and Birney, 2008; Chikhi et al., 2016) (Figure 1C). Compaction yields a graph with locally optimal resolution: regions of the genome which are conserved across individuals are represented by long unitigs, while regions which are highly variable are fractioned into shorter unitigs (Supplementary Figure S1).

We perform GWAS on strains encoded by their unitig (rather than kmer) content, and use the cDBG neighbourhood of significantly associated unitigs as a proxy for their genomic environment. Figure 4 summarises the main steps of the process. The code implementing this process is available at https://gitlab.com/leoisl/dbgwas/ under the GNU Affero General Public License.

**Figure 4:**
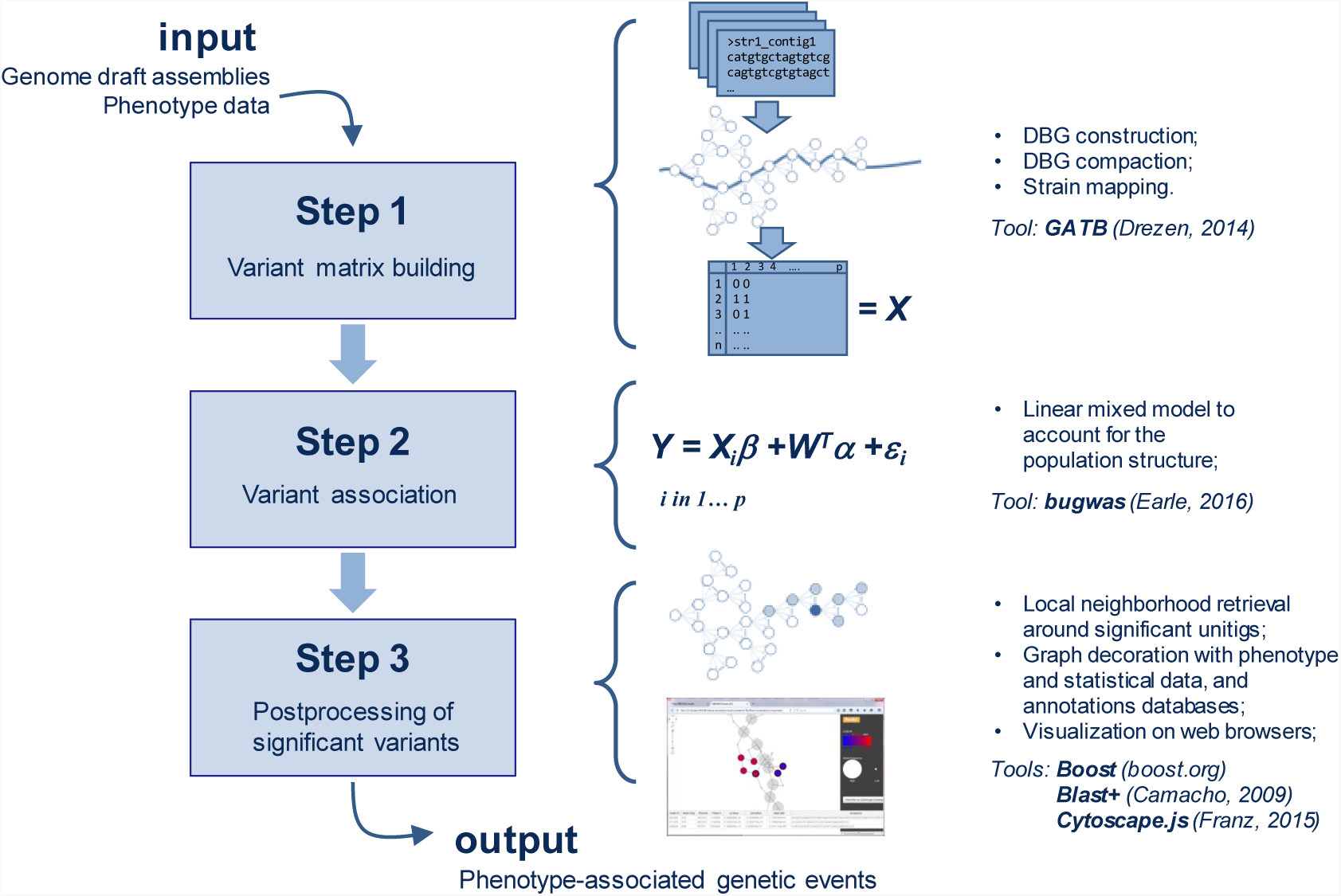
DBGWAS pipeline. DBGWAS takes as input draft assemblies and phenotype data for a panel of bacterial strains. Variant matrix *X* is built in step 1 using cDBG nodes. Variants are tested in step 2 using a linear mixed model. Significant variants are post-processed in step 3 to provide an interactive interface assisting with their interpretation.

### Representing strains by their unitig content (step 1)

#### cDBG construction

We build a single DBG from all genomes given as input using the GATB C++ library (Drezen et al., 2014). We start from contigs rather than reads to be robust to sequencing errors. Consequently, we do not need to filter out low abundance kmers, allowing for the exploration of any variation present in the set of input genomes.

We use a *k* = 31 length for our kmers, as it produced the best performance to retrieve known markers in a pilot experiment (Supplementary Figure S8). The ideal choice of *k*, however, depends on many factors, including the assembly quality, complexity of the input genomes, or presence of repeats. Sensibility analysis to the choice of *k* is extensively presented in Supplementary Section 5. We then compact the DBG using a graph traversal algorithm, which identifies all linear paths in the DBG – each forming a unitig in the cDBG. During this step, we also associate each kmer index to its corresponding unitig index in the cDBG.

#### Unitig presence across genomes

Each genome is represented by a vector of presence/absence of each unitig in the cDBG. To do so, we query the unitig associated to each kmer in a given genome. This procedure is efficient because it relies on constant time operations. Firstly, we use GATB’s Minimal Perfect Hash Function (MPHF) (Limasset et al., 2017) to retrieve the index of a given kmer, and then we use the association between kmer and unitig indexes to know which unitigs the given genome contains. Since these two operations take constant time, producing this vector representation for a genome takes linear time on the size of the genome. It is important to note that the GATB’s MPHF can be successfully applied here because we always use the same list of kmers, *i.e.*, after building the DBG, the set of kmers is fixed and not updated, and because we always query kmers that are guaranteed to be in the DBG (since we do not filter out any kmer).

The unitig description on all the input genomes is stored into a matrix *U*:

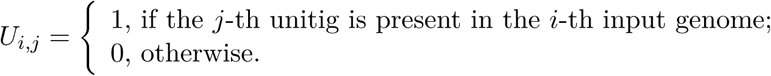

We then transform the matrix *U* into *Z*, giving minor allele description (Earle et al., 2016). *Z* is identical to *U* except for columns with a mean larger than 0.5, which are complemented: *Z*_*j*_ = 1 − *U*_*j*_ for these columns.

We then restrict *Z* to its set of unique columns. If several unitigs have the same minor allele presence pattern, then they will be represented by a single column. Keeping duplicates would lead to performing the same statistical test several times. Finally, we filter out columns whose average is below 0.01. We denote the de-duplicated, filtered matrix of patterns by *X*.

### Testing unitigs for association with the phenotype (step 2)

Human GWAS literature extensively discusses how testing procedures can result in spurious associations if the effect of the population structure is not taken into account (Balding, 2006; Zhou and Stephens, 2014; Widmer et al., 2014). Population structures can be strong in bacteria because of their clonality (Falush and Bowden, 2006; Earle et al., 2016; Lees et al., 2016). A preliminary performance analysis comparing several models for population structure on both simulated and real data (Jaillard et al., 2017a) showed that correcting for population structure using LMMs is often preferable to using a fixed effect correction or not correcting at all.

We thus rely on the bugwas method (Earle et al., 2016), which uses the linear mixed model (LMM) implemented in the GEMMA library (Zhou and Stephens, 2012) to test for association with phenotypes while correcting for the population structure. This method also offers the possibility to test for lineage effects, by calculating p-values for association between the columns of the matrix representing the population structure, and the phenotype (Earle et al., 2016).

Formally, the LMM represents the distribution of the binarized phenotype *Y*_*i*_, given the *j*-th minor allele pattern *X*_*ij*_ and the population structure represented by a set of factors 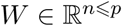, by:

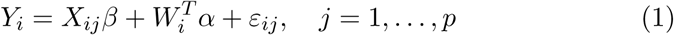

*β* is the fixed effect of the tested candidate on the phenotype, *α* is the random effect of the population structure, and 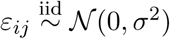 are the residuals with variance *σ*^2^ > 0. *W* is estimated from the *Z* matrix which includes duplicate columns representing both core and accessory genome.

We test *H*_0_: *β* = 0 *versus H*_1_: *β* ≠ 0 in equation 1 for each unitig using a likelihood ratio procedure producing p-values and maximum likelihood estimates 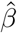. Finally, we compute the q-values, which are the Benjamini-Hochberg transformed p-values controlling for false discovery rate (FDR) in the situation of multiple testing (Benjamini and Hochberg, 1995).

### Interpretation of significant unitigs (step 3)

The LMM can be used to identify deduplicated minor allele presence patterns significantly associated with the phenotype at a chosen level. Because of the deduplication procedure used to build the matrix *X*, each of these patterns can correspond to several unitigs. We now show how the cDBG can be used in the interpretation step.

#### Significance threshold

We select the most significantly associated patterns by defining a Significant Features Filter (*SFF*). In our experiments, we choose not to apply a fixed FDR threshold – even though DBGWAS offers this option, by using a *SFF* value between 0 and 1. Different datasets lead to different q-values, even by several orders of magnitude, and a single FDR threshold would lead to selecting a large number of unitigs generating *>* 1000 subgraphs on some of them (e.g. *S. aureus* ciprofloxacin) as shown in Supplementary Table S8. Instead, we use *SFF* = 100, *i.e.*, retaining the 100 patterns with lowest q-values. However arbitrary, this choice is tractable for all datasets and provides satisfactory results in our experiments. It does not guarantee control of the FDR: only the q-value provides an estimation of the proportion of false discoveries incurred when considering patterns below this value. Checking the q-values of the selected unitigs is therefore essential to assess its significance.

#### Graph neighbourhoods

We define the neighbourhood of each significant unitig *u* (defined by the *SFF*) as the set of unitigs whose shortest path to *u* has at most 5 edges. The objects returned by DBGWAS are the connected components of the graph induced by the neighbourhoods of all significant unitigs in the cDBG. As illustrated in Figure 5, nearby significant unitigs might belong to the same connected component, so this process groups unitigs which are likely to be located closely in the genomes. We refer to the connected components as *subgraphs* in the Results section.

**Figure 5:**
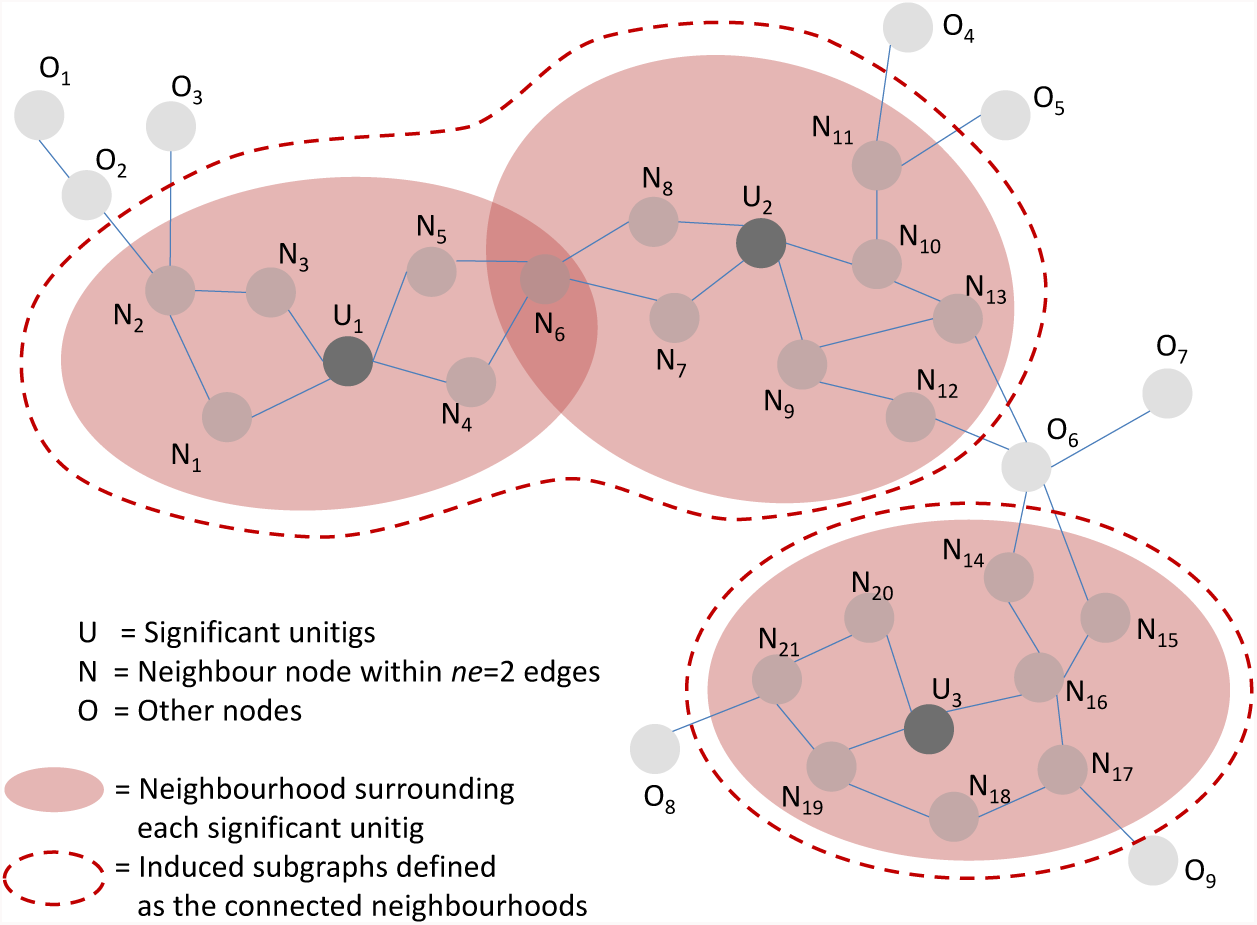
Subgraphs induced by the neighbourhood of significantly associated unitigs. In this example, a neighbourhood of size 2 was used: any unitig distant up to 2 edges from a significant unitig is retrieved to define its neighbourhood. Neighbourhoods are merged if they share at least one node, e.g. the neighbourhoods of *U*_1_ and *U*_2_ are merged because they share *N*_6_, and will be represented in a single subgraph.

The *SFF* value can be tuned to optimise the number and size of the output subgraphs (Supplementary Section 4). The *SFF* value has no impact on subgraphs mostly describing SNPs in core sequences (Supplementary Figures S2). When significant unitigs map to different regions of a single MGE such as a plasmid, several subgraphs are generated but can be gathered into a single subgraph by increasing the *SFF* threshold (Supplementary Figures S4). When signicant unitigs map to several distinct mobile regions which can be found in different contexts (transposon, integron, etc.) at the population level, the resulting subgraph can be huge and highly branching: decreasing the *SFF* threshold allows to select the few most significant unitigs generating a subgraph focusing on the most relevant region (Supplementary Figure S6).

#### Representing metadata with coloured DBGs

The subgraphs are enriched with metadata to make their interpretation easier. We use the node size to represent allele frequencies, *i.e.*, the proportion of genomes containing the unitig sequence. We describe the effect 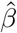 of each unitig as estimated by the LMM using colours, in the spirit of the coloured DBG (Iqbal et al., 2012). Colours are continuously interpolated between red for unitigs with a strong positive effect and blue for those with a strong negative effect.

#### Annotating the subgraphs

DBGWAS offers an optional annotation step using the Blast suite (Camacho et al., 2009) (version 2.6.0+) on local user-defined protein or nucleic acid sequence databases. We annotate the subgraphs of interest by blasting each unitig sequence to the available databases. Users can then easily retrieve the annotations which are the most supported by the nodes in the subgraph, or with the lowest E-value. We provide on the DBGWAS website a resistance determinant database built by merging the ResFinder, MEGARes, and ARG-ANNOT databases (Zankari et al., 2012; Lakin et al., 2017; Gupta et al., 2014), and a subset of UniProt restricted to bacterial proteins (UniProt consortium, 2017). Subgraphs discussed in the Results section were annotated using these databases.

#### Interactive visualization

DBGWAS produces an interactive view of the enriched and annotated subgraphs, allowing the user to explore the graph topology together with information on each node: allele and phenotype frequencies, q-value, estimated effect, and annotation. The view is built using HTML, CSS, and several Javascript libraries, the main one being Cytoscape.js (Franz et al., 2015). Results can be shared and visualized in a web browser. A large number of components can be produced in one run of DBGWAS. We thus provide a summary page allowing the user to preview and filter the subgraphs. Filtering can be based upon the minimum q-value of all unitigs in the component (min_*q*_), or based on the annotations. A complete description of the DBGWAS interactive interface is available in Supplementary Section 6.

### Datasets

We used in our experiments genome sequences from three bacterial species with various degrees of genome plasticity, from more clonal to more plastic: *Mycobacterium tuberculosis*, *Staphylococcus aureus*, and *Pseudomonas aeruginosa*. We build a fourth panel (see below WHO list panel), used only for time and memory performance assessment and defined according to the top-3 WHO priority pathogens list^1^. These panels are summarised in Table 2.

**Table 2:** **Panels used in this study.** We selected 3 bacterial species for their distinctly differing levels of genome plasticity, plus an inter-species panel integrating the top-3 WHO priority pathogens list.

| Panel name | Species | Genome plasticity | Range of genome length | Source |
| --- | --- | --- | --- | --- |
| TB | *M. tuberculosis* | very low | 4.4 Mbp | (Davis et al., 2016) |
| SA | *S. aureus* | low | 2.7-3.1 Mbp | (Gordon et al., 2014) |
| PA | *P. aeruginosa* | high | 5.8-7.6 Mbp | (van Belkum et al., 2015) |
| WHO list | *A. baumannii*<br>*P. aeruginosa*<br>*K. pneumoniae*<br>*E. coli*<br>*Enterobacter sp.*<br>*E. cloacae* | high | 3.5-7.6 Mbp | PATRIC |

#### TB panel

*M. tuberculosis* (TB) is a human pathogen causing 1.7 million deaths each year^2^. This species is known for its apparent absence of horizontal gene transfer (HGT) and accordingly, most of the reported resistance determinants are chromosomal mutations (Gygli et al., 2017) in core genes or gene promoters. Intergenic regions are also described to be instrumental in multidrug-resistance (MDR) and extensively drug-resistant (XDR) phenotypes (Zhang et al., 2013). We use the PATRIC AMR phenotype data, as well as genome assemblies from their resource (Wattam et al., 2016; Davis et al., 2016). We thus gather a total of 1302 genomes after filtering based on genome length. Phenotype data include isoniazid, rifampicin, streptomycin, ethambutol, ofloxacin, kanamycin and ethionamide resistance status. Except for the last three drugs, phenotype data are available for more than a thousand genomes. We reconstruct MDR and XDR phenotypes based on the WHO definition^3^. XDR phenotype could only be defined for 689/1302 strains as it required data for at least 4 drugs. Information on how phenotype data and genome assemblies were obtained is available on the PATRIC website.

#### SA panel

*S. aureus* is a human pathogen causing life-threatening infections. It is subject to HGT and many plasmids, mobile elements, and phage sequences have been described in its genome. However, this does not affect the species’ genome size which is always close to 3 Mbp (Mlynarczyk et al., 1998). Most antibiotic resistance mechanisms are well determined by known variants as shown in a previous study (Gordon et al., 2014). This study obtained an overall sensitivity of 97% for predicting 12 phenotypes from rules based on antibiotic marker mapping. We use this study panel of 992 strains obtained by merging their derivation and validation sets.

#### PA panel

*P. aeruginosa* is a ubiquitous bacterial species responsible for various types of infections. It is highly adaptable thanks to its ability to exchange genetic material within the species. The species accessory genome is particularly important both in terms of size and diversity and carries more than half of the genetic determinants already described to confer resistance to antimicrobial drugs (Kung et al., 2010; van Belkum et al., 2015; Jaillard et al., 2017b). We use a panel of 282 strains, gathered from two collections which mostly include clinical strains: the bioMérieux collection (van Belkum et al., 2015) (*n*=219) and the Pirnay collection (Pirnay et al., 2009) (*n*=63). Genome assemblies and categorical phenotypes for 9 antibiotics are available (Jaillard et al., 2017b). Binarised phenotypes of amikacin resistance are available on the DBGWAS project page to provide this dataset as an example for users.

#### WHO list panel

This panel is built from PATRIC AMR Phenotype data and genome resource and is designed to search for resistance determinants which are shared by the top-3 pathogens in the WHO priority list, all Gram negative: *Acinetobacter baumannii* carbapenem-resistant, *P. aeruginosa* carbapenem-resistant, and Enterobacteriaceae carbapenem-resistant, ESBL-producing.

We collate all genomes having a phenotype for at least one of the antibiotics belonging to the carbapenem family (imipenem, meropenem, ertapenem or doripenem). It represents 234 genomes with phenotype data for *A. baumannii*, 125 for *P. aeruginosa*, 135 for *K. pneumoniae*, 6 for *E. coli*, 3 for *Enterobacter sp.*, and 2 for *E. cloacae*.

#### Phenotype binarisation

Most available phenotypes are categorical, with S, I and R levels, respectively, for susceptible, intermediary, and resistant. We binarise them by assigning a zero value to susceptible strains (S) and one to others (I and R).

### Resistome-based GWAS (RWAS)

RWAS are performed to validate that DBGWAS retrieves all known determinants found by a targeted approach. In this validation study we used bugwas with the same phenotypes and population structure matrix *W* so the RWAS analyses and DBGWAS only differ by their input variant matrix (unitigs *versus* SNPs or genes presence/absence).

#### P. aeruginosa

We use the variant matrix described previously (Jaillard et al., 2017b), which includes presence/absence of known resistance genes and gene variants, as well as all SNPs called against a reference sequence of these genes (and gene variants).

#### M. tuberculosis

We build the variant matrix using the same approach as for *P. aeruginosa* (Jaillard et al., 2017b): we call the SNPs from a list of known resistance genes (Coll et al., 2015; Gygli et al., 2017; Palomino and Martin, 2014) (available in Supplementary Section 3.1).

We sort the rows of the output file by q-values. Tables S6 and S7 summarise all top variants using their q-value ranks, while Figure 3 reports the annotations of all variant with a q-value *<* 0.05 for *M. tuberculosis* streptomycin and *P. aeruginosa* levofloxacin resistance.

### Kmer-based GWAS

We benchmarked DBGWAS, SEER (Lees et al., 2016) and HAWK (Rahman et al., 2017) in terms of computational efficiency (running time and memory usage), simplicity of use and downstream analyses (Supplementary Section3.2), and the ability to retrieve known markers (see Figure 3).

#### SEER

We installed SEER static precompiled v1.1.3. SEER’s pipeline is mainly composed of four steps: 1) Kmer counting; 2) Population structure estimation; 3) Running SEER; 4) Downstream analysis. For running these steps with the correct parameters, we followed the tutorial available on SEER’s github page: for kmer counting, we used fsm-lite and for step 2, we used Mash v2.0 (Ondov et al., 2016). In step 3, we used a `--maf 0.01`. Downstream analysis involved getting the kmers that were called significant by SEER, sorting them by LRT p-value, blasting them against the two databases presented in Section *step 3*, keeping the best hit for each kmer.

#### HAWK

We installed HAWK v0.8.3-beta. HAWK’s pipeline comprises five steps: 1) Kmer counting; 2) Running HAWK; 3) Assembling significant kmers; 4) Getting statistics on the assembled sequences; 5) Downstream analysis. The first four steps were performed as described in HAWK’s github page. However, in the first step, we had to remove the lower-count cutoff in `jellyfish` dump (parameter -L), since we are working with contigs and not reads. Moreover, for assembling the significant kmers, we used ABYSS v2.0.2 (Jackman et al., 2017). Finally, the last step was performed similarly as the one described for SEER.

## Data and source code access

All data used in this work were previously published.

Data generated by our method and discussed in the manuscript are available at http://leoisl.gitlab.io/DBGWAS_support/experiments.

The source code and precompiled version of our method is available on gitlab: https://gitlab.com/leoisl/dbgwas/.

## Acknowledgments

The authors thank Jean-Baptiste Veyrieras, Sarah Earle, Chieh-Hsi Wu and Daniel Wilson, as well as Jean-Pierre Flandrois, Manolo Gouy, Stéphane Schicklin and Ghislaine Guigon for their insightful comments. LL acknowledges CNPq/Brazil for the financial support. This work was performed using the computing facilities of the CC LBBE/PRABI. LL is funded by the Brazilian Ministry of Science, Technology and Innovation (in portuguese, Ministério da Ciência, Tecnologia e Inovação - MCTI) through the National Counsel of Technological and Scientific Development (in portuguese, Conselho Nacional de Desenvolvimento Científico e Tecnológico - CNPq), under the Science Without Borders (in portuguese, Ciências Sem Fronteiras) scholarship grant process number 203362/2014-4. VL is funded by the Agence Nationale de la Recherche ANR-12-BS02-0008 (Colib’read) and ANR-16-CE23-0001 (ASTER). LJ is funded by the Agence Nationale de la Recherche ANR-14-CE23-0003-01 (MACARON) and ANR-17-CE23-0011-01 (FAST-BIG).

## Author contributions

MJ and LJ designed the method with the help of VL and MT. LL, LJ and MJ implemented the code available on gitlab. MJ, LL and PM ran the experiments described in this paper. MJ, LL, LJ, PM and AvB wrote the manuscript. All authors have reviewed and approved the final version of the manuscript.

## Footnotes

1 http://www.who.int/mediacentre/news/releases/2017/bacteria-antibiotics-needed/en/

2 http://www.who.int/mediacentre/factsheets/fs104/en/

3 http://www.who.int/tb/areas-of-work/drug-resistant-tb

